# Selfish bacteria are active throughout the water column of the ocean

**DOI:** 10.1101/2021.07.26.453833

**Authors:** Greta Giljan, Sarah Brown, C. Chad Lloyd, Sherif Ghobrial, Rudolf Amann, Carol Arnosti

## Abstract

Heterotrophic bacteria use extracellular enzymes to hydrolyze high molecular weight (HMW) organic matter to low molecular weight (LMW) hydrolysis products that can be taken into the cell. These enzymes represent a considerable investment of carbon, nitrogen, and energy, yet the return on this investment is uncertain, since hydrolysis of a HMW substrate outside a cell yields LMW products that can be lost to diffusion and taken up by scavengers that do not produce extracellular enzymes^1^. However, an additional strategy of HMW organic matter utilization, ‘selfish’ uptake^2^, is used for polysaccharide degradation, and has recently been found to be widespread among bacterial communities in surface ocean waters^3^. During selfish uptake, polysaccharides are bound at the cell surface, initially hydrolyzed, and transported into the periplasmic space without loss of hydrolysis products^2^, thereby retaining hydrolysate for the selfish bacteria and reducing availability of LMW substrates to scavenging bacteria. Here we show that selfish bacteria are common not only in the sunlit upper ocean, where polysaccharides are freshly produced by phytoplankton, but also deeper in the oceanic water column, including in bottom waters at depths of more than 5,500 meters. Thus, the return on investment, and therefore also the supply of suitable polysaccharides, must be sufficient to maintain these organisms.

---

High molecular weight carbohydrates – polysaccharides - constitute a major fraction of both living and detrital marine organic matter^4,5^. The degradation of polysaccharides depends largely on the activities of bacteria equipped with the extracellular enzymes required to dismantle these often highly complex structures to low molecular weight (LMW) pieces (e.g., ^6^). Since LMW hydrolysis products may also be taken up by ‘scavengers’ that do not produce the enzymes, the enzyme ‘producers’ that carry out external hydrolysis might benefit only in part from their own enzyme activities^1^. Selfish bacteria circumvent this problem by transporting large polysaccharide fragments into their periplasmic space, minimizing loss of hydrolysis products^2^. A broad range of polysaccharides is taken up via selfish mechanisms by diverse bacteria in surface ocean waters^7^; the speed and extent of selfish uptake and external hydrolysis vary by geographic location^7,8^, as well as by the nature and abundance of polysaccharides at different phytoplankton bloom stages^9^. The extent to which selfish bacteria are present and active in other depths of the ocean, however, remains unexplored.

Given that polysaccharide-hydrolyzing enzymes are exquisitely specific for substrate structure^10^, we hypothesized that selfish bacteria would be most abundant and active in locations and depths at which freshly produced – structurally unaltered – polysaccharides are common. Since both HMW dissolved organic matter and particulate organic matter are more abundant and are ‘fresher’ – have a higher fraction of chemically characterizable components – in the upper ocean than in the deep ocean^11,12^, we expected that selfish bacteria would be particularly dominant in the upper water column. We therefore collected water samples at three stations characterized by different physical, chemical, and productivity conditions in the western North Atlantic: in the Gulf Stream, in productive waters off of the coast of Newfoundland, and in the oligotrophic waters of the North Atlantic Gyre (Fig. S1). At these stations, we collected water from the surface, deep chlorophyll maximum (33 to 104 meters), upper mesopelagic (∼300 m), and bottom (3190 to 5580 m). Triplicate incubations were made with water from three different Niskin bottles from each depth (biological replicates). We quantified the presence and activity of selfish bacteria by adding small quantities of structurally-distinct fluorescently-labeled polysaccharides (FLA-PS) and incubating these water samples at *in situ* temperatures. The added FLA-PS – laminarin, pullulan, fucoidan, xylan, chondroitin sulfate, and arabinogalactan – have different monomer compositions and linkage types. These polysaccharides were chosen because they are abundant in marine algae and phytoplankton and/or because a wide range of marine bacteria may produce enzymes that hydrolyze them (e.g., ^13-16^). We concurrently measured selfish uptake and external hydrolysis rates of the FLA-PS, quantified cell abundances, measured bacterial protein production, and tracked bacterial community composition.

Much to our surprise, selfish bacteria were abundant at all water depths that we investigated. These bacteria were identified microscopically by the co-localization of the blue DAPI staining of DNA and the associated intense green staining from the FLA-PS (Fig. 1). Considerable selfish activity was evident even at the t0 timepoint, when 14-17% of bacteria in surface water, 5-22% at the DCM, 5-8% at 300 m, and 5-12% of bacteria in bottom water took up one of the FLA-PS (Fig. 2a-d). With increasing incubation time, the proportion of cells taking up one of the FLA-PS increased, especially in sub-surface waters. Selfish uptake reached a maximum of 13-18% of DAPI-stainable cells in surface waters, 14-26% at the DCM, 12-18% at 300 m, and 25-67% in bottom water. Uptake at the t0 timepoint reflects the short-term response of the *in situ* community, since the time elapsed between substrate addition and sample processing is likely insufficient for major changes in community composition, while later timepoints reflect the activities of a community that has changed in composition with time.

**Figure 1.**
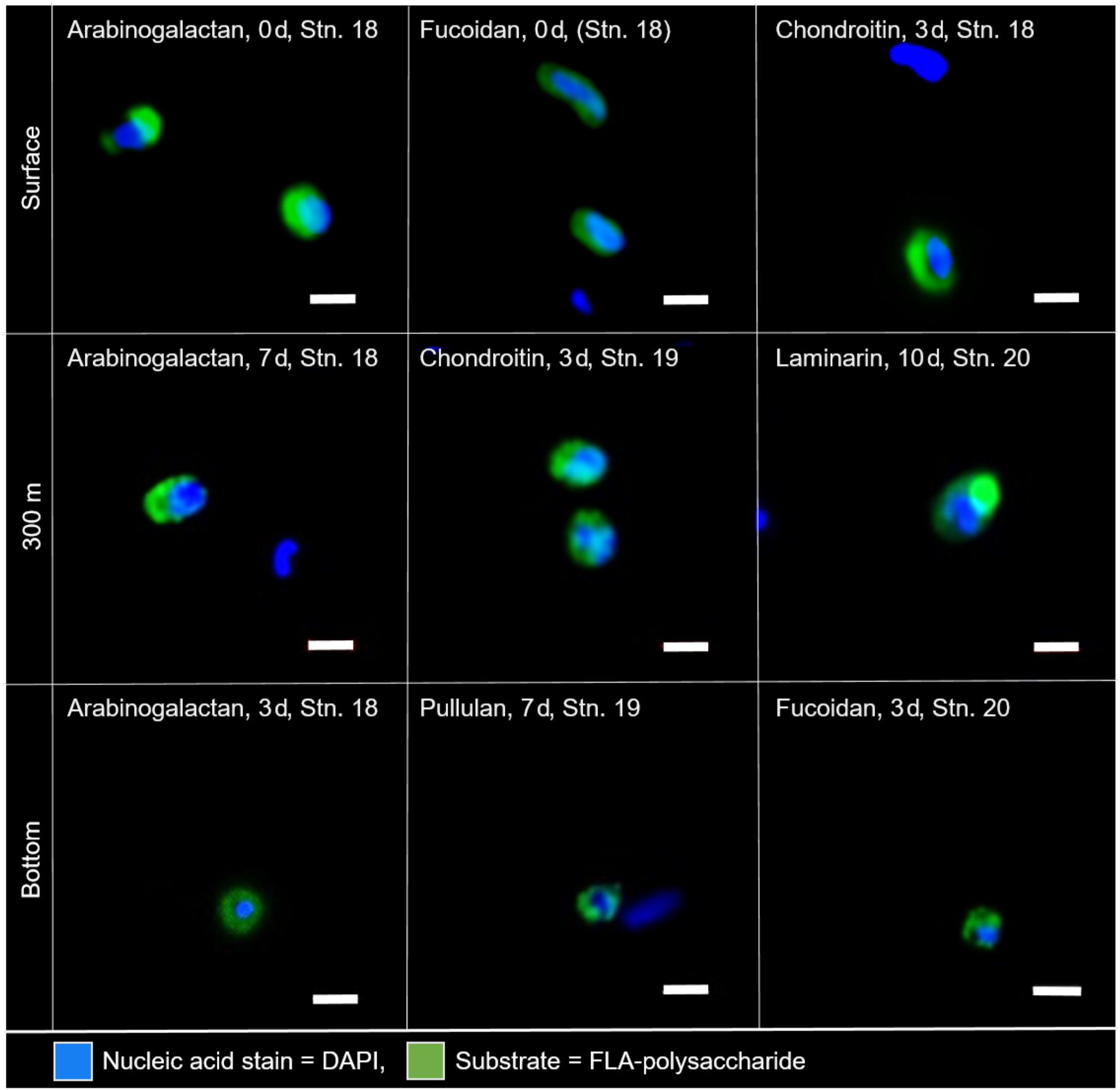
Selfish bacteria throughout the water column in the western North Atlantic Ocean. Super-resolution images of microbial cells from surface water, 300 m water depth, and bottom water (3,190 m, 4,325 m, and 5,580 m, respectively) showing accumulation of fluorescently labeled arabinogalactan, fucoidan, chondroitin sulfate, laminarin, and pullulan due to selfish uptake. Blue signal (DAPI) shows nucleic acids, green signal is due to fluoresceinamine-labeled polysaccharides. Scale bar = 1 µm.

**Figure 2.**
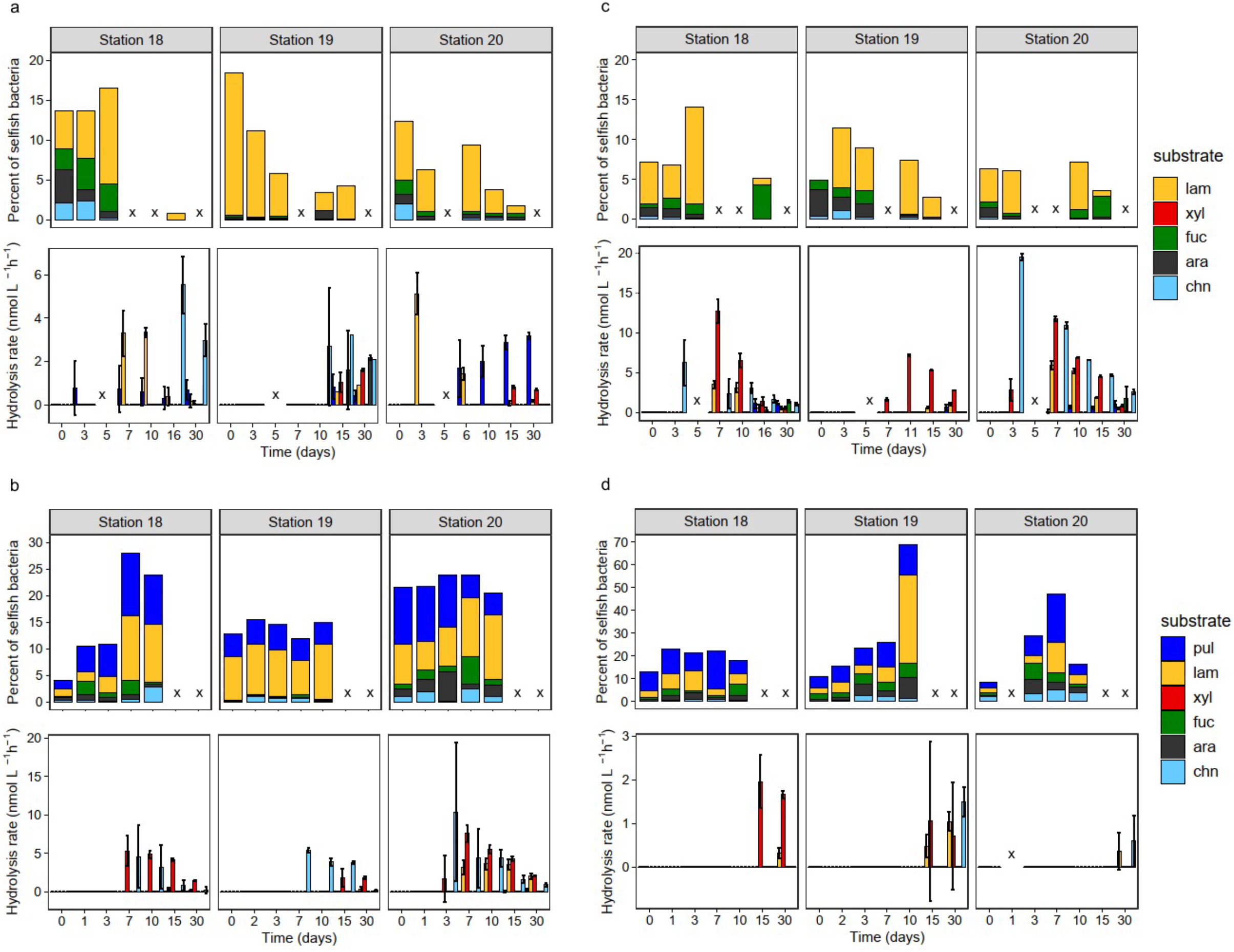
Heterotrophic polysaccharide utilization throughout the water column at three distinct locations in the western North Atlantic Ocean. Selfish uptake and extracellular (external) hydrolysis of six different fluorescently labeled polysaccharides (FLA-PS) in **(a)** surface waters, **(b)** at the DCM, **(c)** at 300 m water depth, and **(d)** in bottom waters at Stations 18, 19 and 20 over the course of individual FLA-PS amended incubations. Note that selfish FLA-xylan uptake at all stations and FLA-pullulan uptake in surface waters and at 300 m depth could not be analyzed due to high background fluorescence and are therefore not included in the data. Error bars represent the average of biological replicates (n = 3). Samples marked with an x were not analyzed.

The *in situ* bacterial communities showed considerable selfish uptake, despite initial station- and depth-related differences in composition (Fig. 3). These initial communities changed markedly for the most part over the time course of incubation (Fig. 4; Figs S2-S5), but selfish uptake at most stations and depths remained constant or increased after the t0 timepoint. The compositional changes in unamended incubations were very similar to the incubations amended with FLA-PS, demonstrating that the addition of FLA-PS by themselves had little influence on community composition (Figs. S2c-S5c) or cell counts (Fig. S6). Moreover, the changes in vcommunity composition typically were not convergent for different depths (Fig. S7), in that different genera dominated even in cases where similar classes became more abundant with time. For example, although Gammaproteobacteria became relatively more abundant with time in many of the incubations (Fig. 4, Figs. S2-S5), the dominant phylotypes varied by depth and station (Figs. S8-S10); there were no obvious connections between specific changes in bacterial community composition and selfish uptake.

**Figure 3.**
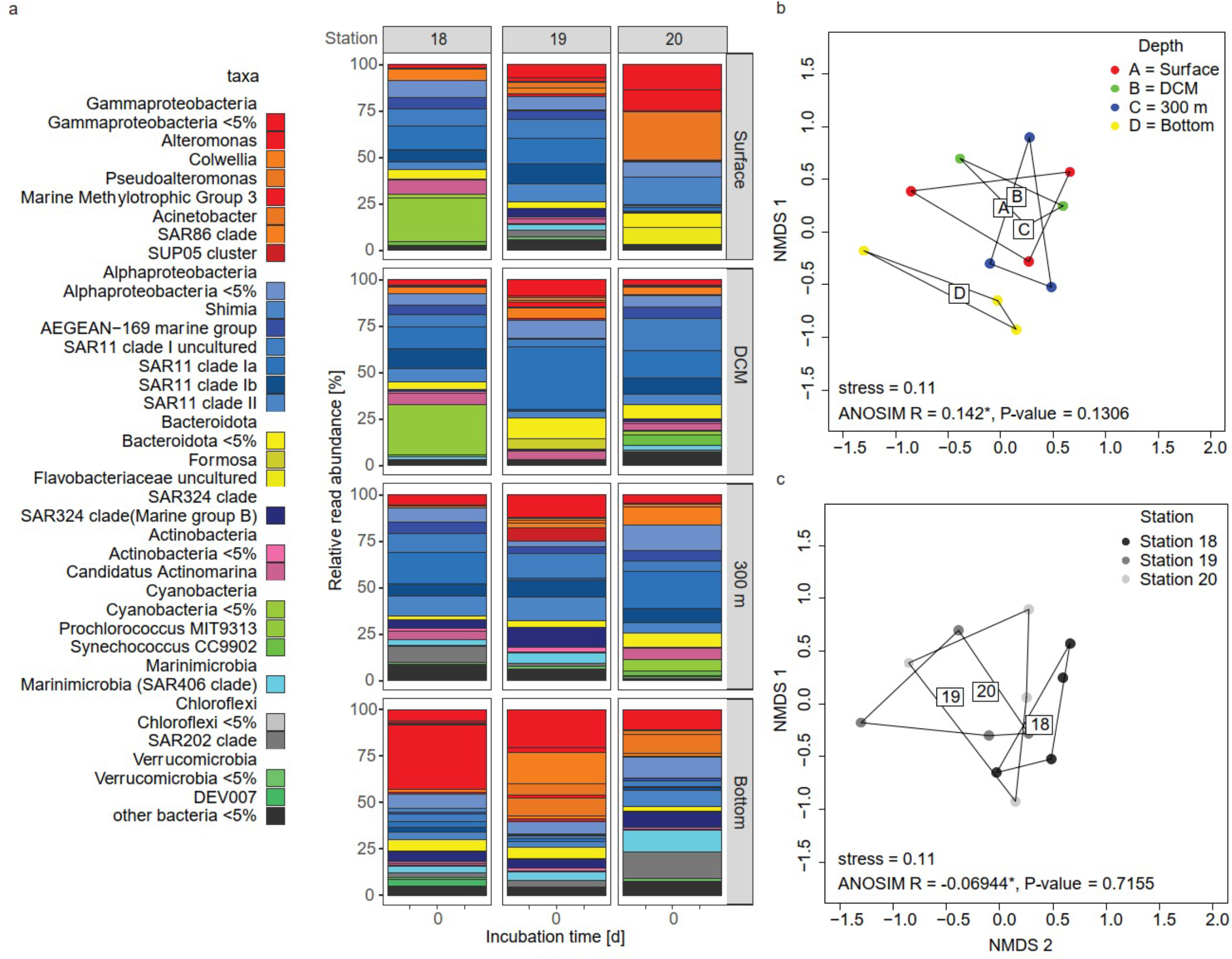
The initial bacterial communities at four depths at three distinct locations in the western North Atlantic. Bacterial communities **(a)** differ more by **(b)** depth but are also different by **(c)** station. * denotes statistically significant results. Each community represents the average of biological replicates (n = 3).

**Figure 4.**
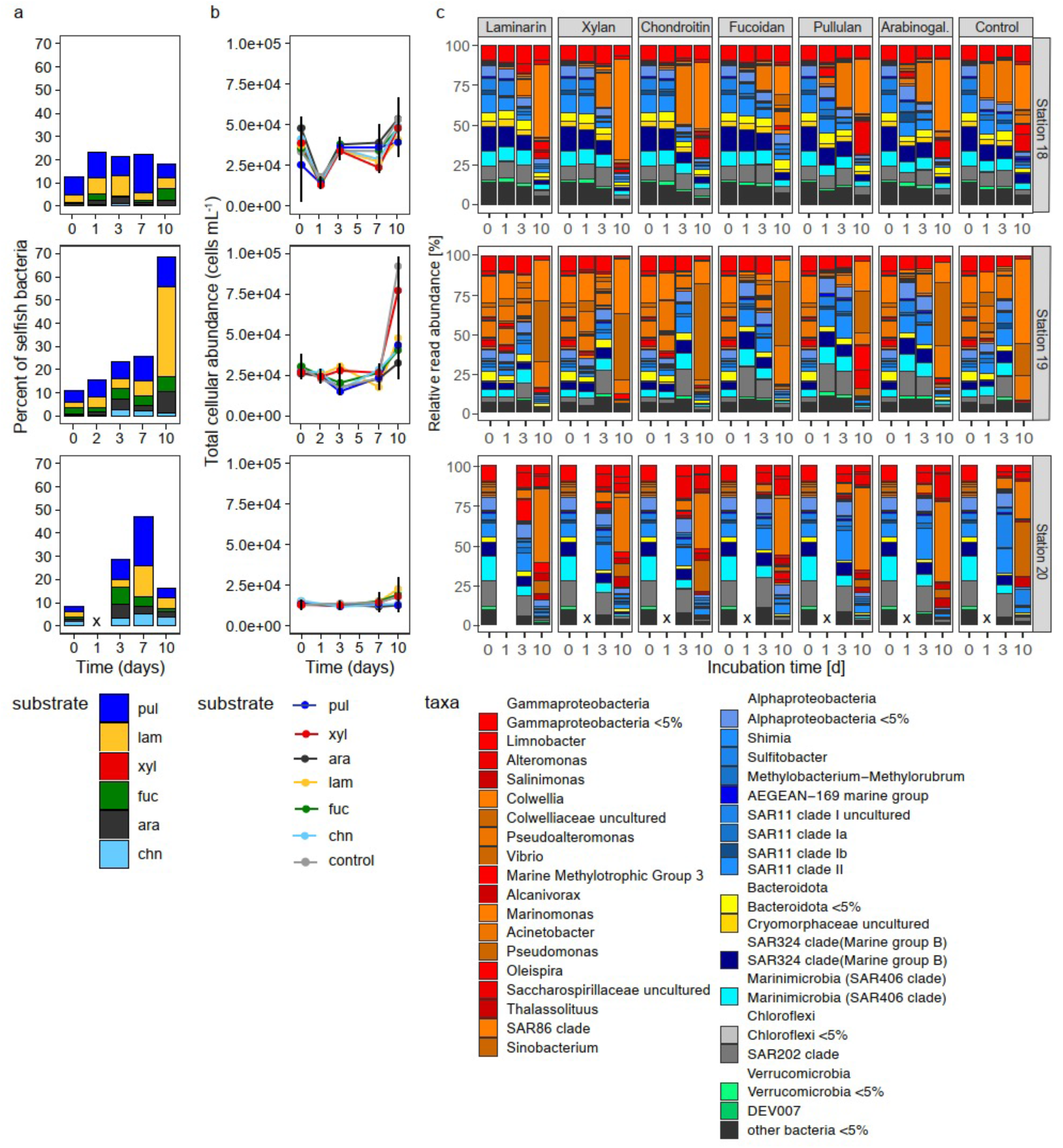
Selfish polysaccharide uptake, cell counts, and bacterial community composition in North Atlantic bottom water. **(a)** Selfish uptake of FLA-PS over the course of 10 day FLA-PS amended incubations at Stations 18, 19 and 20 (same data as in Fig. 2d). **(b)** Development of the total microbial cell counts and **(c)** bacterial community composition in each of the FLA-PS amended incubations from (a), and in the unamended treatment control. The initial community shortly after the addition of the FLA-PS is depicted at 0 days of incubation. Bars represent the average of up to 3 replicates. Time points marked with an x were not analyzed.

Although selfish uptake mechanisms have been studied most intensively in members of the gut-dwelling Bacteroidetes^17,18^, our previous investigations in surface ocean waters demonstrated that a range of bacteria, including members of the Bacteroidetes, Planctomycetes, Verrucomicrobia, and the genus *Catenovulum* (Gammaproteobacteria) carry out selfish uptake^7,19^. Furthermore, a large fraction of the selfish population is still unidentified^7,19^. The observation of selfish activity at multiple depths throughout the water column against a background of changing bacterial community composition suggests that this strategy of substrate acquisition is widespread, and in the ocean is not restricted to a limited range of bacterial taxa. Since selfish uptake cannot be inferred solely from genomic information^20^, however, we cannot yet determine whether selfish uptake mechanisms are as widely distributed as the ability to produce extracellular enzymes to carry out external hydrolysis.

We observed remarkable differences in patterns of selfish uptake and external hydrolysis, which were measured in the same incubations. While selfish uptake was measurable for a broad range of polysaccharides at all stations and all depths, external hydrolysis was more variable among stations at similar depths, and decreased sharply in bottom water compared to surficial waters (Fig. 2). In surface waters at Stn. 19, for example, all polysaccharides except fucoidan were externally hydrolyzed, but at Stns. 18 and 20 only laminarin, chondroitin, and xylan were externally hydrolyzed; these spatial variations in external hydrolysis are consistent with previous observations in surface ocean waters^21^. Station-related variability was also evident at a depth of 300 m: at Stns. 18 and 20, a broad spectrum of polysaccharides was hydrolyzed, but at Stn. 19, only xylan, laminarin, and pullulan were hydrolyzed. In bottom waters of all three stations, only two or three polysaccharides were hydrolyzed – laminarin and xylan at Stn. 18, laminarin, xylan, and chondroitin at Stn. 19, and laminarin and chondroitin at Stn. 20 – at comparatively low rates, first evident at late incubation timepoints. Overall, the spectrum of substrates hydrolyzed was narrower and the hydrolysis rates in deep water were considerably lower than in surface water, consistent with the few previous reports of polysaccharide hydrolysis in deep ocean waters^22-24^. Patterns of external hydrolysis and selfish uptake at the same stations and depths thus showed striking contrasts.

The presence of selfish bacteria in the water column, also at depths well below the euphotic zone, requires new consideration of the economics of substrate processing and uptake. Two-player models of enzyme producers and scavengers have considered the conditions under which extracellular enzyme production may pay off (e.g.,^1^), such as when polysaccharides are abundant^25^, or when they are found in sufficiently dense patches^26^. Inclusion of selfish bacteria in this calculus, as described in a new conceptual model^27^, suggests that substrate structural complexity, as well as abundance, needs to be taken into account. Selfish uptake, in which hydrolysate is efficiently captured, can help ensure that investment in extracellular enzymes generates sufficient return. From this perspective, selfish uptake appears to be widespread i) when a substrate is highly complex and requires considerable enzymatic investment, or ii) when there is high competition for a very widely-available substrate, such that competition is a primary consideration.

The first case covers enzymatic investment to acquire comparatively rare and structurally complex substrates that are selfishly taken up, as in the deep ocean (Fig. 4). Arguably, intact polysaccharides are likely a comparatively rare commodity in most of the subsurface ocean, and thus should be a target for selfish uptake, a consideration that would explain the broad range of substrates selfishly taken up in much of the water column. Polysaccharides such as fucoidan are particularly good examples for a potential pay-off from selfish uptake, since fucoidan hydrolysis requires an extraordinary investment in extracellular enzymes^28^. Moreover, external hydrolysis of fucoidan is comparatively rarely detected in the surface ocean^21^, and to date has not been detected in deep ocean waters^29,22-24^. The second set of circumstances for which selfish uptake pays off – high competition for an abundant polysaccharide – applies especially to laminarin. Oceanic production of laminarin has been estimated to be on the order of 12-18 gigatons annually^13,30^, providing a vast supply of readily-degradable substrate to heterotrophic microbial communities. Moreover, external hydrolysis of laminarin is measurable in almost every site and location in the ocean investigated to date^21, 27, 22-24, 3, 7-9^, pointing at extraordinarily widespread capabilities to utilize this polysaccharide. Selfish uptake of this polysaccharide therefore ensures return on enzyme investment by capturing a substrate that would otherwise be acquired by competitors.

Here we show for the first time that selfish uptake occurs at multiple depths in the water column, including in deep bottom waters. Furthermore, rapid selfish uptake of polysaccharides in bottom water provides important clues about the physiology of bacteria in the deep ocean, as well as the nature of the substrates that they use. Approximately 5-10% of the bacterial community at our three deep ocean sites was ready and able to take up specific polysaccharides shortly after addition (Figs. 1-2, Fig. 4). This response is notable in light of the observation that uptake of a simple amino acid at these depths (as demonstrated by leucine used for bacterial productivity measurements; Table S1), which does not require prior enzymatic hydrolysis, was quite low especially at Stns. 18 and 19. These observations together suggest that a bacterial strategy focused on rapid uptake of structurally more-complex, higher molecular weight polysaccharides pays off in deep water because there is a sufficient supply of these substrates, whereas the *in situ* inventory of individual amino acids in bottom water is likely too low^31^ to merit special targeting by bacteria. Although we currently lack data on the polysaccharide component of POM in bottom waters, a new method to specifically quantify laminarin in POM has demonstrated a considerable laminarin concentration in POM in the upper water column (including measurements to a depth of 300m^30^). Moreover, time-variable rapid transport of bacteria/particles to bottom water depths has been demonstrated^32^; some of this organic matter reaching these depths evidently is fresh, and has not been thoroughly worked over in the upper ocean; this organic matter could include intact polysaccharides. Recent measurements of DOM in deep ocean waters additionally suggest that high molecular weight polysaccharides are added to the DOM pool circulating in the deep ocean^33^.

Evidence of selfish uptake of highly complex polysaccharides in the deep ocean supports the point that the flux of relatively unaltered organic matter to the deep ocean must be of sufficient magnitude^34-35^ to support a viable and reactive population of selfish bacteria in the deep. The presence of bacteria that are capable of processing complex substrates in a selfish manner also points out that measurements of bacterial metabolism that are dependent upon uptake of monomeric substances have likely underestimated an important fraction of heterotrophic carbon cycling activity, since the enzymatic systems used for selfish uptake are specifically tuned to their target substrates^2^. The prevalence of selfish uptake against a backdrop of changing bacterial community composition (Fig. 2, Fig. 4, Figs. S2-S5), moreover, suggests that selfish uptake as a substrate acquisition strategy pays off sufficiently that it is comparatively widespread among bacteria. Selfish uptake is important not only in the surface waters of the ocean, but also in the upper mesopelagic and deep ocean, and is carried out by heterotrophic bacteria whose carbon cycling activities help drive much of the marine carbon cycle.

## Methods

### i. Station location and seawater collection

Seawater was collected at three stations in the western North Atlantic aboard the research vessel *Endeavor* (cruise EN638) using a sampling rosette of 30-liter Niskin bottles fitted with a Sea-Bird 32 conductivity-temperature-depth (CTD) profiler, between May 15^th^ and 30^th^ 2019 (Fig. S1). Collection depths included surface water (2.5–6 m water depth), the deep-chlorophyll maximum (DCM; depth identified via chorophyll fluorescence signal of the CTD: 104 m, 33 m, 64 m water depth at Stns. 18, 19, and 20, respectively), ∼300 m (300 m at Stns. 18 and 20; 318 m at Stn. 19), and bottom water (3,190 m, 4,325 m, and 5,580 m, at Stns. 18, 19, and 20, respectively; Fig. S1).

At each station and depth, triplicates of 600 mL (DCM and bottom water) or 290 mL (surface and 300 m water) were added to sterile, acid rinsed (10% HCl) bottles and incubated for up to 30 days in the dark at *in situ* temperatures (room temperature (RT) for water from the surface, DCM and 300 m; 4°C for bottom water) with one of the six FLA-PS: arabinogalactan, chondroitin sulfate, fucoidan, laminarin, pullulan and xylan, each at 3.5 µM monomer equivalent concentration. A single live treatment control without the addition of any substrate was included for the DCM and bottom waters; autoclaved killed controls were included for each substrate at each station and each depth, and were incubated under the same conditions alongside polysaccharide incubations

Subsamples for microbial cell counts and selfish FLA-PS uptake were collected from DCM and bottom water incubations 0, 1, 3, 7, and 10 days after the addition of polysaccharides; in surface and 300 m incubations, subsamples were collected 0, 3, 7, 10, and 15 days after polysaccharide addition. Note also that the t0 timepoint measurements of selfish uptake represent a time period of ca. 30 min (surface, 300m) to 5 hrs (DCM, bottom water), representing the time required after initial substrate addition to go back and process all of the samples to which substrate had been added. To measure the extracellular hydrolysis of FLA-PS, subsamples were collected on days 0, 3, 7, 10, 15, and 30 of the incubations. Subsamples for bulk community analysis were taken before the addition of FLA-PS and at day 1, 3, and 10 of the incubation with DCM and bottom water and at day 0, 3, 7, 10, and 15 in the surface and 300 m incubations.

### ii. Synthesis of FLA-PS and measurements of extracellular enzymatic activities

Arabinogalactan, chondroitin sulfate, fucoidan, laminarin, pullulan, and xylan were fluorescently labeled with fluoresceinamine (Sigma) and characterized according to Arnosti (2003)^36^. Subsamples (2 ml) were removed at days 0, 1, 3, 7, 10, 15, and 30 days post FLA-PS addition, and analyzed after Arnosti (2003)^36^. Note that the added substrate is in competition with naturally occurring substrates, and thus calculated hydrolysis rates are potential hydrolysis rates.

### iii. Counts of total and substrate-stained cells

#### Cell counts

To prepare samples, 25-50 mL of 1% FA fixed sample were filtered onto a 0.22 µm pore size polycarbonate filter at a maximum vacuum of 200 mbar. The DNA of filtered cells was counterstained using 4′,6-diamidin-2-phenylindol (DAPI) and mounted with a Citifluor/VectaShield (4:1) solution. A minimum of 45 microscopic images per sample were aquired as described by Bennke *et al*. (2016)^37^ with a fully automated epifluorescence microscope (Zeiss AxioImager.Z2 microscope stand, Carl Zeiss) equipped with a cooled charged-coupled-device (CCD) camera (AxioCam MRm + Colibri LED light source, Carl Zeiss), three light-emitting diodes (UV-emitting LED, 365 nm for DAPI; blue-emitting LED, 470 nm for FLA-PS 488) and a HE-62 multi filter module with a triple emission filter (425/50 nm, 527/54 nm, LP 615 nm, including a triple beam splitter of 395/495/610, Carl Zeiss) using a 63x magnification oil immersion plan apochromatic objective with a numerical aperture of 1.4 (Carl Zeiss). Final cell enumeration on the acquired images was performed using the image analysis software ACMETOOL (http://www.technobiology.ch and Max Planck Institute for Marine Microbiology, Bremen). Automated cell counts were checked manually.

Total microbial cell numbers and FLA-PS stained cells were counted in a single experimental setup, following Reintjes et al. (2017)^3^. Counting validation was done through manual cell counting on all stains. Selfish substrate uptake could be measured for only four or five of the six polysaccharides used; at all stations and depths, xylan incubations yielded high background fluorescence, which interfered with cell counting; this problem also affected efforts to count cells for pullulan uptake in surface waters and at a depth of 300 m. Note also that we report the fraction of cells carrying out selfish uptake under the assumption that each substrate is taken up by different bacteria (i.e., when reporting that for example 22% of total DAPI-stainable cells were substrate-stained, we add together the percentages taking up laminarin, fucoidan, arabinogalactan, and chondroitin). Since selfish uptake of each substrate is measured in different incubations (triplicate incubations of each individual substrate), however, it is possible that some or all of the cells taking up one substrate also take up another substrate via a selfish mechanism.

### iv. Super-resolution imaging of selfish polysaccharide uptake

The specific substrate accumulation pattern in FLA-PS stained cells was visualized on a Zeiss LSM780 with Airyscan (Carl Zeiss) using a 405 nm, a 488 nm, and a 561 nm laser with detection windows of 420-480 nm, 500-550 nm, and LP 605 nm, respectively. Z-stack images of the cells were taken with a Plan-Apochromat 63x/1.4 oil objective and the ZEN software package (Carl Zeiss) was used for subsequent AiryScan analysis.

### v. Taxonomic bacterial community analysis

The initial bacterial communities and their change over the course of the incubation were determined through bulk 16S rRNA analysis. Therefore, 25 mL samples from each incubation were filtered onto a 0.22 µm pore size polycarbonate filter at a maximum vacuum of 200 mbar, dried and frozen at -20 °C until further processing. Total DNA extraction from filter was done using the DNeasy Power Water Kit (Quiagen). Determination of the concentration as well as the size of the extracted DNA was done via gel chromatography using a Fragment Analyzer™ (Advanced Analytical). Amplification of the variable 16S rRNA regions V3 and V4 (490 bp) was done in 30 cycles using the 5 PRIME HotMasterMix (Quantabio) together with the Bakt_314F (CCTACGGGNGGCWGCAG) and Bakt_805R (GACTACGVGGGTATCTAATCC)^38^ PCR primer pair with an individual 8 bp barcode adapter (based on the NEB Multiplex Oligos for Illumina, New England Biolabs) attached to the forward primer and the reverse primer. The amplified PCR product was purified and size selected using the AMPure XP PCR Cleanup system (Beckman Coulter). Barcoded products were pooled in equimolar concentrations and send for paired-end Illumina sequencing (2×250 bp HiSeq2500) to the Max Planck-Genome-center Cologne. Sequences were merged, demultiplexed and quality trimmed (sequence length 300–500 bp, < 2% homopolymers, < 2 % ambiguities) with BBTools^39^. The SILVAngs pipeline^40^ with the SSU rRNA SILVA database 138 was used for sequence comparison and taxonomic assignment of the retrieved sequences.

### vi. Statistical analysis of bacterial communities

Analysis of the bacterial community composition was done normalized reads representing > 1,000 reads per sample and the average of triplicates for the FLA-PS amended incubations was used for further analysis and visualization. Archaeal and eukaryal reads were excluded from analysis. Differences in the community composition between station, water depth, incubation time, and substrate amended to unamended incubations were analyzed by analysis of similarity (ANOSIM) and visualized in non-metric multi-dimensional scaling (NMDS) plots, using Bray-Curtis dissimilarity matrices. The community shift over the course of the incubation was visualized by the comparison of the read abundance on genus level from the initial community to the respective read abundance in the incubations over time.

### vii. Bacterial productivity

Bacterial productivity was measured after Kirchman et al. (2001)^41^. In brief, bacterial protein production was calculated from leucine incorporation rates, measured in samples that were incubated at *in-situ* temperatures in the dark for time periods of 12 to 24 h. Bacterial carbon production was calculated by multiplying bacterial protein production by 0.86^42,41^.

### viii. Data availability

Bacterial 16S rRNA gene sequences were archived as Illumina-generated libraries at the European Nucleotide Archive (ENA) of The European Bioinformatics Institute (EMBL-EBI) under the accession number PRJEB45894.

## Acknowledgments

We thank the captain and crew of R/V *Endeavor*, as well as the other members of the scientific party of the EN638 cruise, for excellent work at sea. Andreas Ellrot (MPI Bremen) provided essential support for the microscopic analyses. This project was funded by NSF OCE-1736772 to CA, with additional funding by the Max Planck Society.

## Author contributions

Conceived and planned project: CA, RA. Carried out work at sea: GG, SB, CCL, SG, RA, CA. Analyzed samples post-cruise: GG, SB, CCL, SG. Created figures: GG, SB. Wrote manuscript: GG, SB, CA with input from all co-authors.

