## Supplemental figures for "Selfish bacteria are active throughout the water column of the ocean"

a

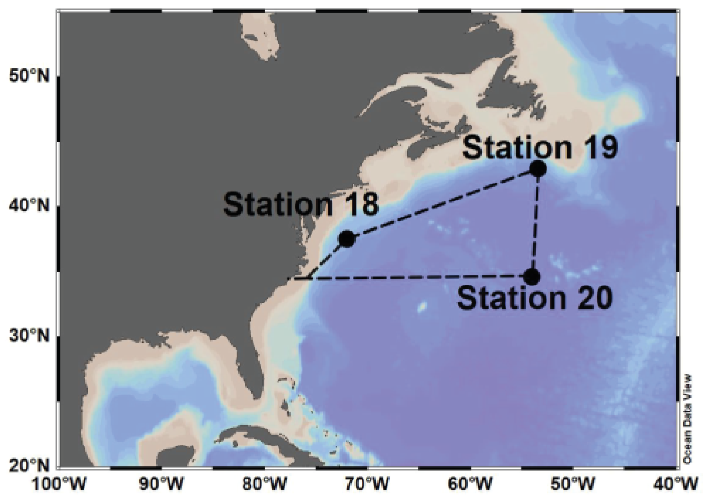


b

| Station | Station 18 | | | |
|---|---|---|---|---|
| Depth name | **Surface** | **DCM** | **300 m** | **Bottom** |
| Depth (m) | 2.5 | 104 | 300 | 3,190 |
| Temperature (°C) | 25.18 | 23.44 | 18.83 | 2.29 |
| Salinity (PSU) | 36.42 | 36.67 | 36.61 | 34.89 |
| Oxygen (mL $L^{-1}$) | 4.48 | 4.53 | 4.27 | 5.89 |
| Chl. Fluor. (mg/$m^3$) | 0.15 | 0.40 | 0.07 | 0.12 |

| Station | Station 19 | | | |
|---|---|---|---|---|
| Depth name | **Surface** | **DCM** | **300 m** | **Bottom** |
| Depth (m) | 6 | 33 | 318 | 4,325 |
| Temperature (°C) | 8.78 | 9.30 | 9.92 | 2.19 |
| Salinity (PSU) | 33.72 | 33.94 | 35.21 | 34.88 |
| Oxygen (mL $L^{-1}$) | 6.63 | 6.45 | 3.34 | 5.73 |
| Chl. Fluor. (mg $m^{-3}$) | 1.48 | 1.33 | 0.13 | 0.06 |

| Station | Station 20 | | | |
|---|---|---|---|---|
| Depth name | **Surface** | **DCM** | **300 m** | **Bottom** |
| Depth (m) | 5 | 64 | 300 | 5,580 |
| Temperature (°C) | 21.03 | 20.09 | 18.31 | 2.18 |
| Salinity (PSU) | 36.73 | 36.68 | 36.56 | 34.85 |
| Oxygen (mL $L^{-1}$) | 4.92 | 5.03 | 4.68 | 5.46 |
| Chl. Fluor. (mg $m^{-3}$) | 0.11 | 0.21 | 0.05 | 0.06 |

c

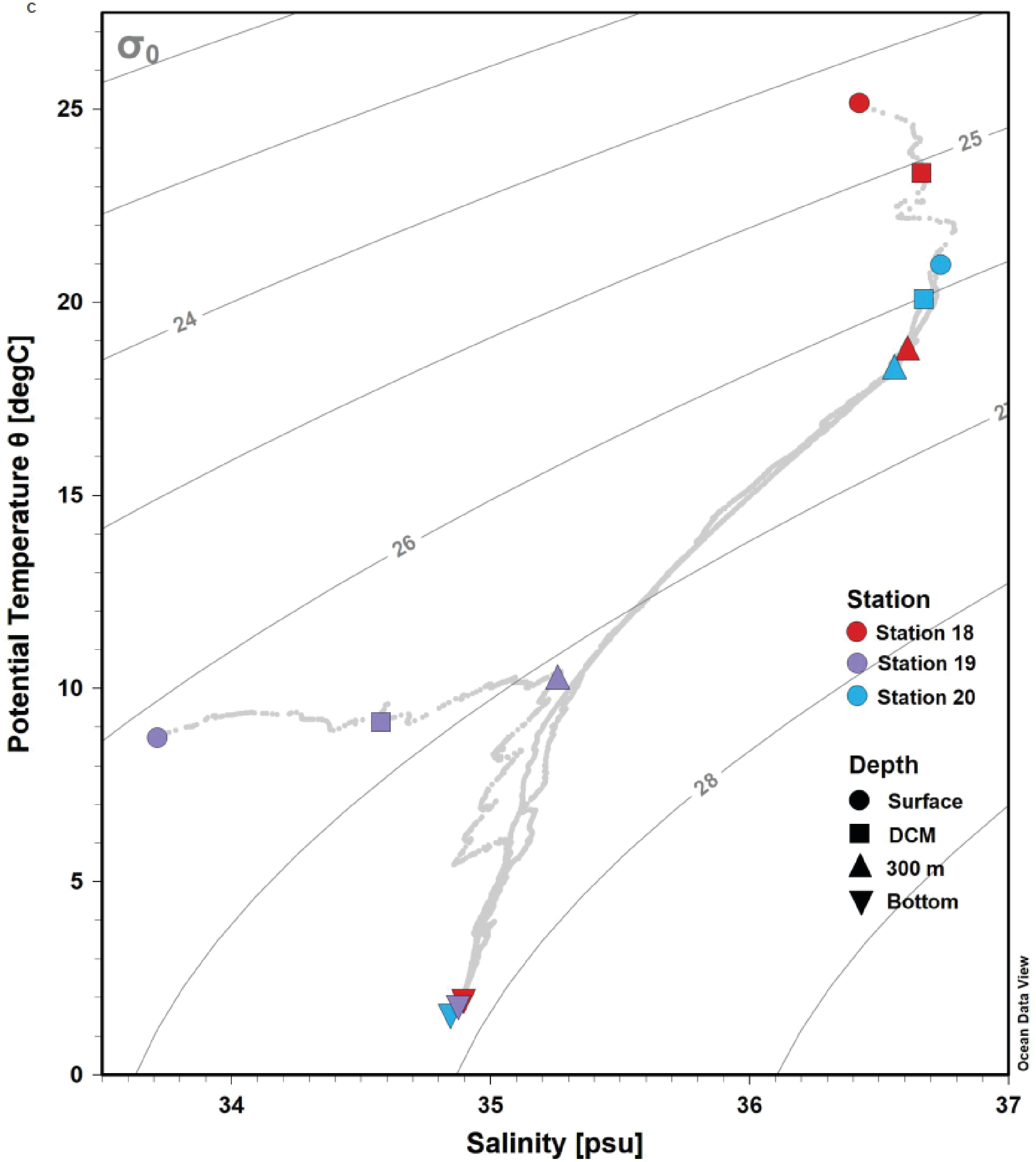


**Figure S1** Chemical and physical station characteristics at Stations 18, 19, and 20 of cruise EN638: **(a)** station locations **(b)** temperature, salinity, dissolved oxygen, and chlorophyll fluorescence and **(d)** T-S plot showing water mass characteristics throughout the water column at each station. Specific depths sampled are showed with symbols (legend).

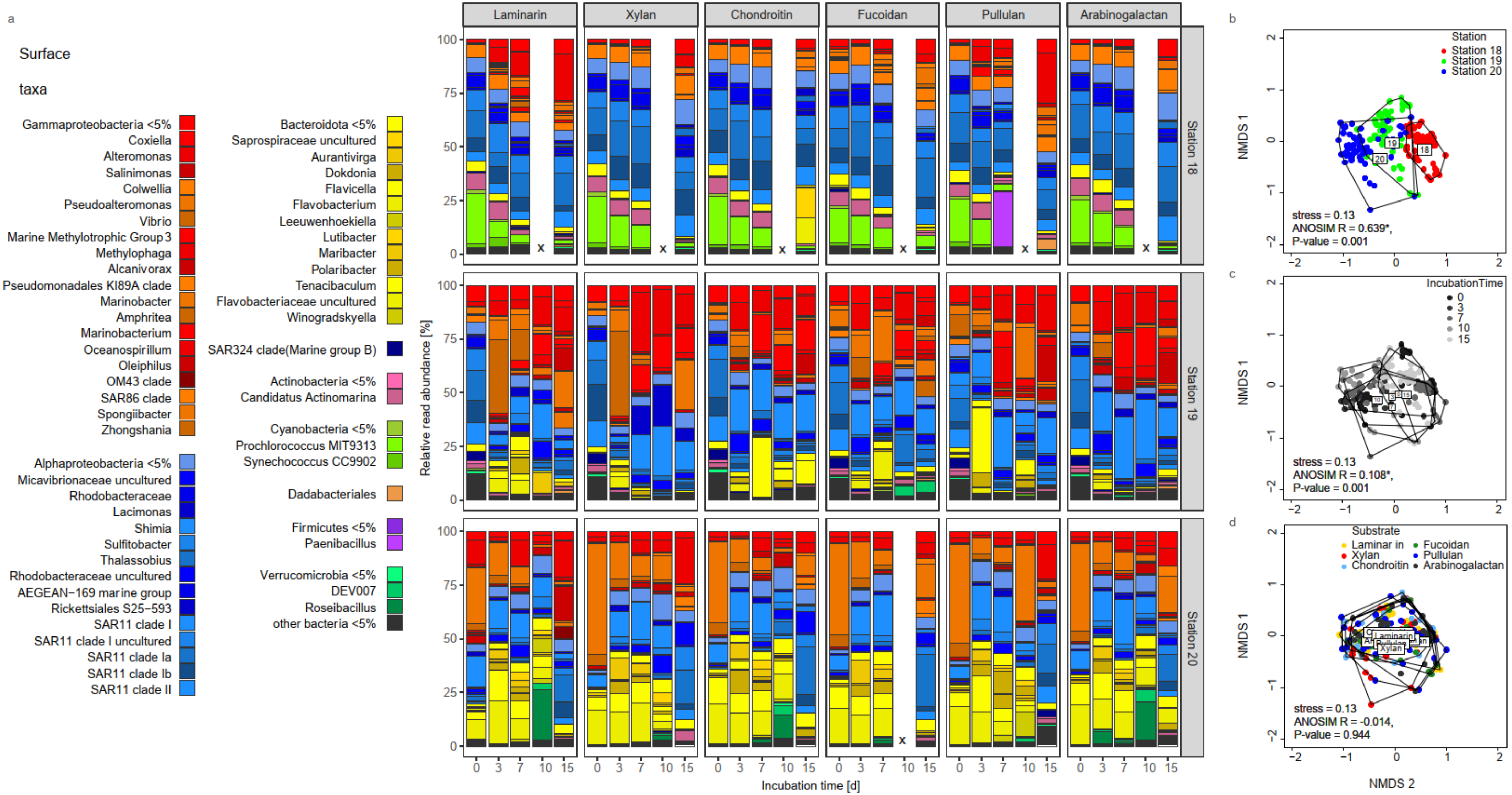


**Figure S2** Changes in bacterial community composition in incubations of surface water from Stns. 18-20, three distinct locations in the western North Atlantic. **(a)** Development of the bacterioplankton community in FLA-laminarin, FLA-xylan and FLA-chondroitin, FLA-pullulan, FLA-arabinogalactan and FLA-fucoidan amended incubations. The initial community shortly after the addition of the FLA-PS is depicted at 0 days of incubation. Bars represent the average of up to 3 replicates. Communities marked with an x were not analyzed. Non-metric multidimensional scaling (NMDS) plot based on Bray-Curtis dissimilarity in bacterial communities clusters the communities by **(b)** station and **(c)** incubation time, but **(d)** not by added polysaccharide. This analysis was confirmed by statistical analysis of similarity (ANOSIM) performed with 99 permutations on the community composition. * denotes statistically significant results.

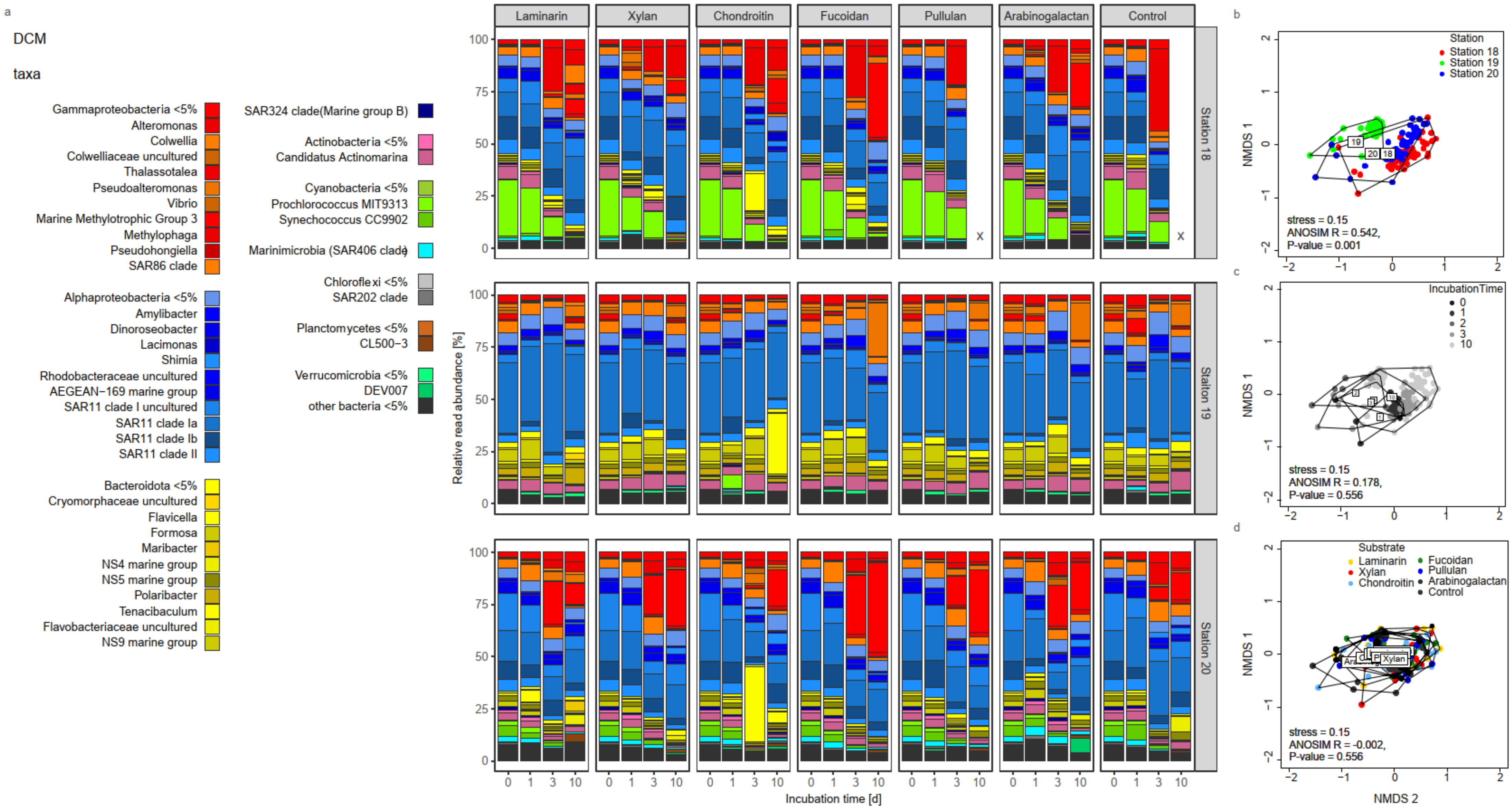


**Figure S3** Changes in bacterial community composition in incubations of water from the deep chlorophyll maximum layer at Stns. 18-20, three distinct locations in the western North Atlantic. **(a)** Development of the bacterioplankton community in FLA-laminarin, FLA-xylan and FLA-chondroitin, FLA-pullulan, FLA-arabinogalactan and FLA-fucoidan amended incubations in comparison to an unamended control. The initial community shortly after the addition of the FLA-PS is depicted at 0 days of incubation. Bars represent the average of up to 3 replicates. Communities marked with an x were not analyzed. Non-metric multidimensional scaling (NMDS) plot based on Bray-Curtis dissimilarity in bacterial communities clusters the communities by **(b)** station and **(c)** incubation time but **(d)** not by added polysaccharide. Results were confirmed by statistical analysis of similarity (ANOSIM) performed with 99 permutations on the community composition. * denotes statistically significant results.

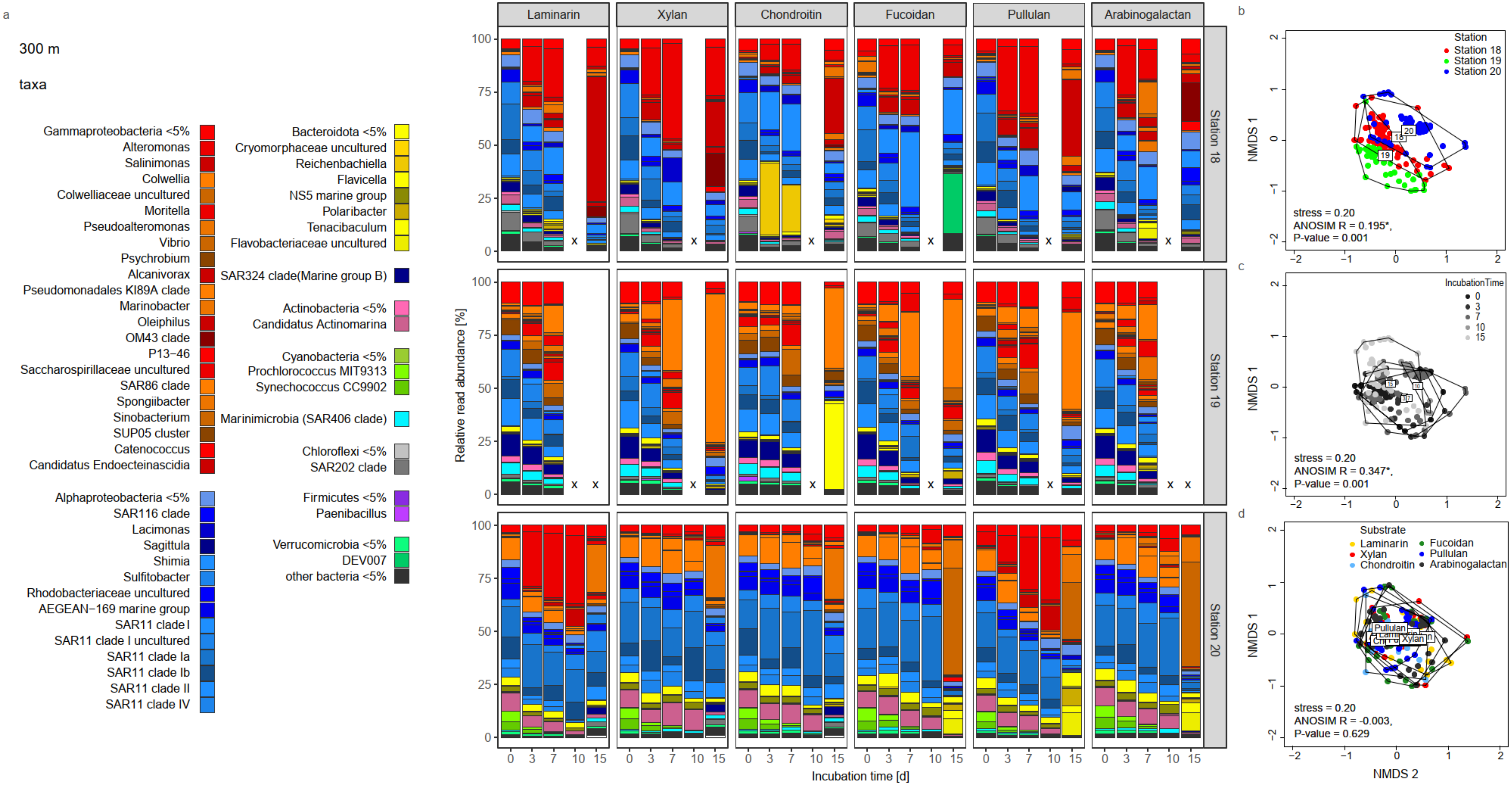


**Figure S4** Changes in bacterial community composition in incubations of water from a depth of 300m at Stns. 18-20, three distinct locations in the western North Atlantic. **(a)** Development of the bacterioplankton community in FLA-laminarin, FLA-xylan and FLA-chondroitin, FLA-pullulan, FLA-arabinogalactan and FLA-fucoidan amended incubations. The initial community shortly after the addition of the FLA-PS is depicted at 0 days of incubation. Bars represent the average of up to 3 replicates. Communities marked with an x were not analyzed. Non-metric multidimensional scaling (NMDS) plot based on Bray-Curtis dissimilarity in bacterial communities clusters the communities by **(b)** station and **(c)** incubation time but **(d)** not by added polysaccharide. These results were confirmed by statistical analysis of similarity (ANOSIM) performed with 99 permutations on the community composition. * denotes statistically significant results.

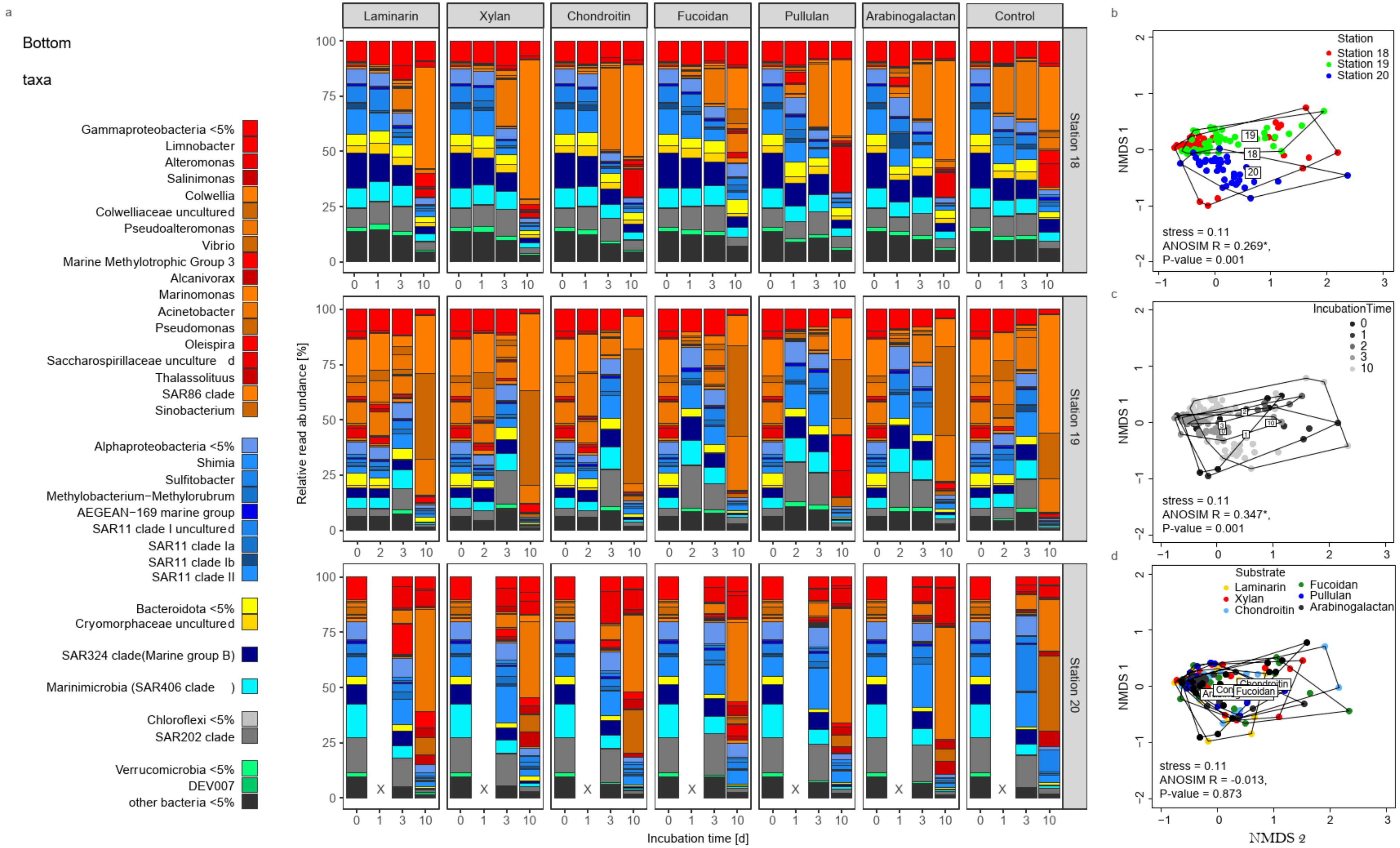


**Figure S5** Changes in bacterial community composition in incubations of bottom water from Stns. 18-20, three distinct locations in the western North Atlantic. Depths were 3,190m, 4,325m, and 5,580m for Stns. 18, 19, and 20, respectively. **(a)** Development of the bacterioplankton community in FLA-laminarin, FLA-xylan and FLA-chondroitin, FLA-pullulan, FLA-arabinogalactan and FLA-fucoidan amended incubations in comparison to an unamended control. The initial community shortly after the addition of the FLA-PS is depicted at 0 days of incubation. Bars represent the average of up to 3 replicates. Communities marked with an x were not analyzed. Non-metric multidimensional scaling (NMDS) plot based on Bray-Curtis dissimilarity in bacterial communities clusters the communities by **(b)** station and **(c)** incubation time but **(d)** not by added polysaccharide. These results were confirmed by statistical analysis of similarity (ANOSIM) performed with 99 permutations on the community composition. * denotes statistically significant results.

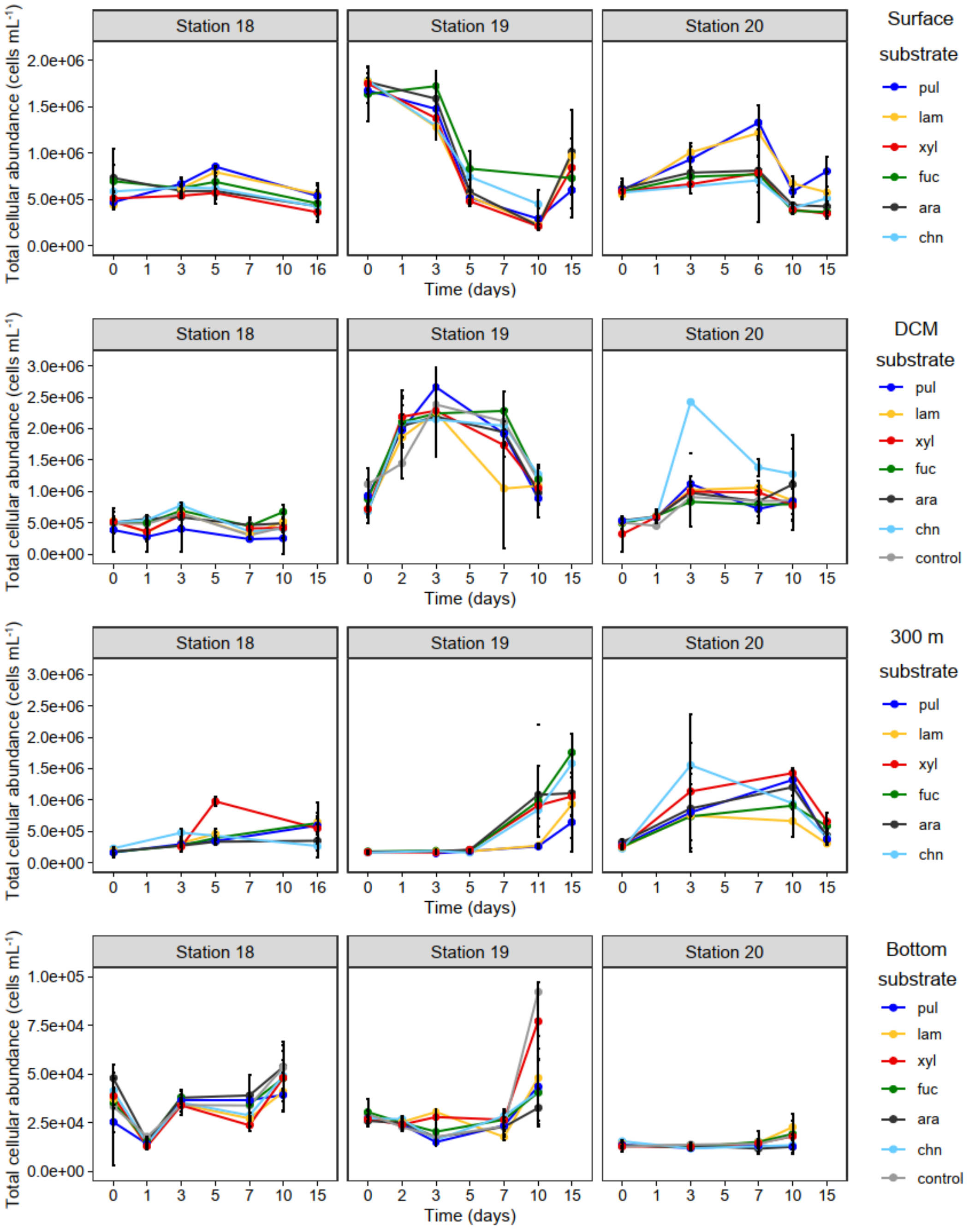


**Figure S6** Development of total microbial cell counts in FLA-PS amended incubations in water from the surface, DCM, 300m depth, and bottom water collected at Stns. 18, 19, and 20. Cell counts are shown over the 15 day time course of incubation with the fluorescently labeled polysaccharides laminarin, xylan, chondroitin sulfate, pullulan, arabinogalactan, and fucoidan, plus an unamended control for incubations from the DCM and in bottom water. Error-bars of the amended incubations represent the average of three replicates (n = 3).

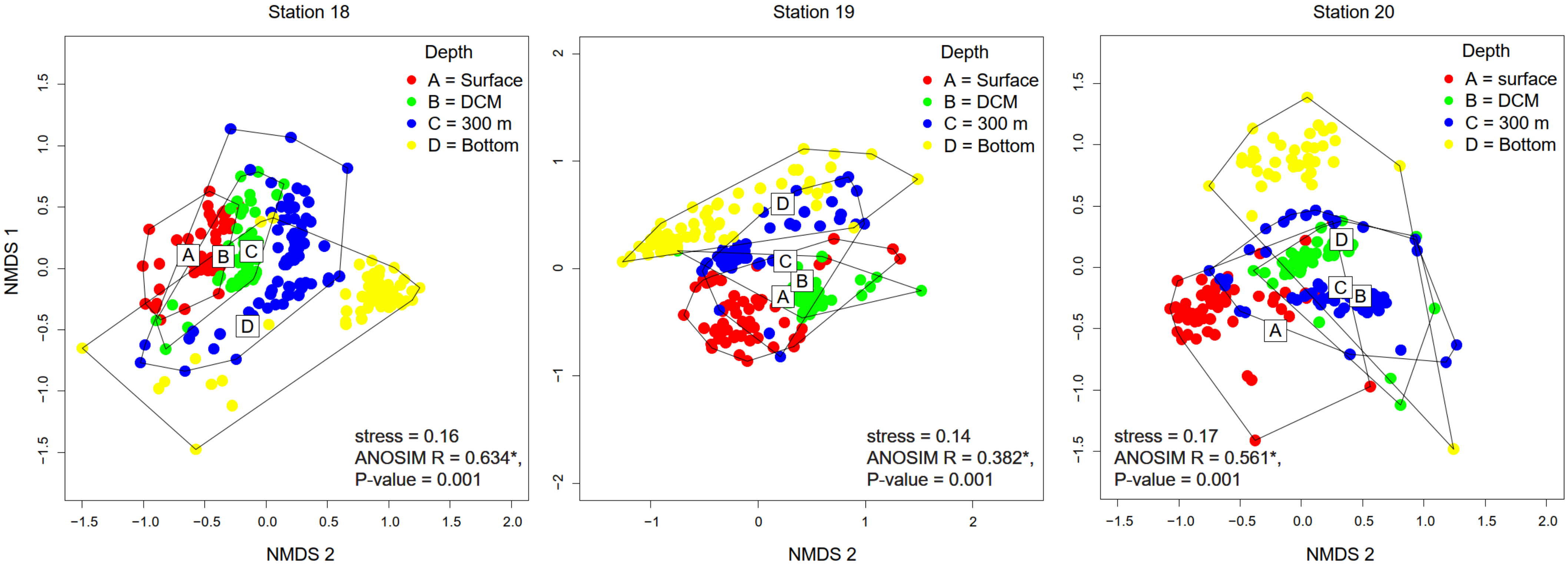


**Figure S7** Non-metric multidimensional scaling (NMDS) plot based on Bray-Curtis dissimilarity demonstrates differences in bacterial communities clustered by depth at Stns. 18 - 20. These results were confirmed by statistical analysis of similarity (ANOSIM) performed with 99 permutations on the community composition. * denotes statistically significant results.

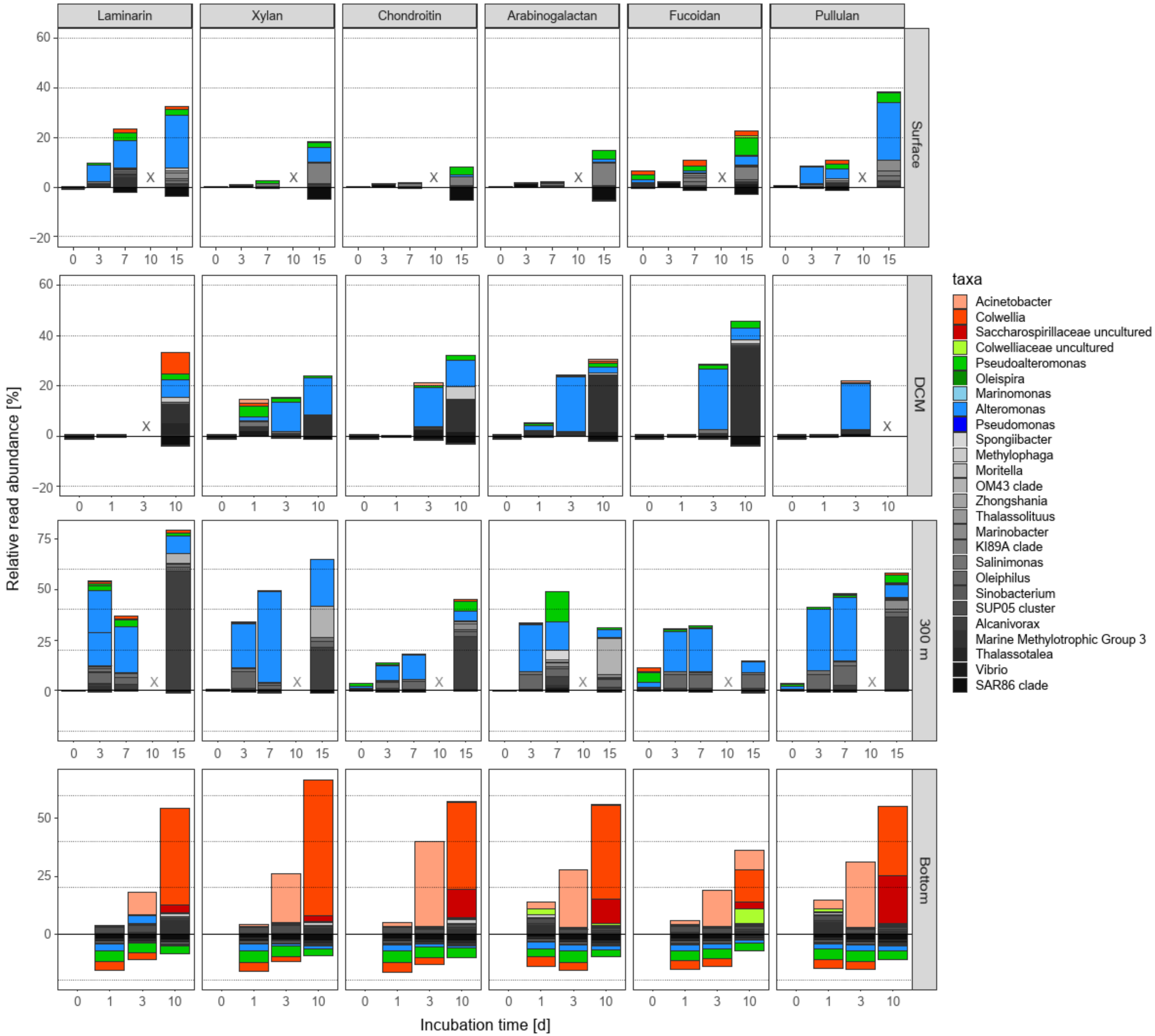


**Figure S8** Development of individual gammaproteobacterial taxa in FLA-PS amended incubations throughout the water column at Station 18. Bars represent the shift in 16S rRNA read abundance of individual Gammaproteobacteria relative to the initial bacterial community.

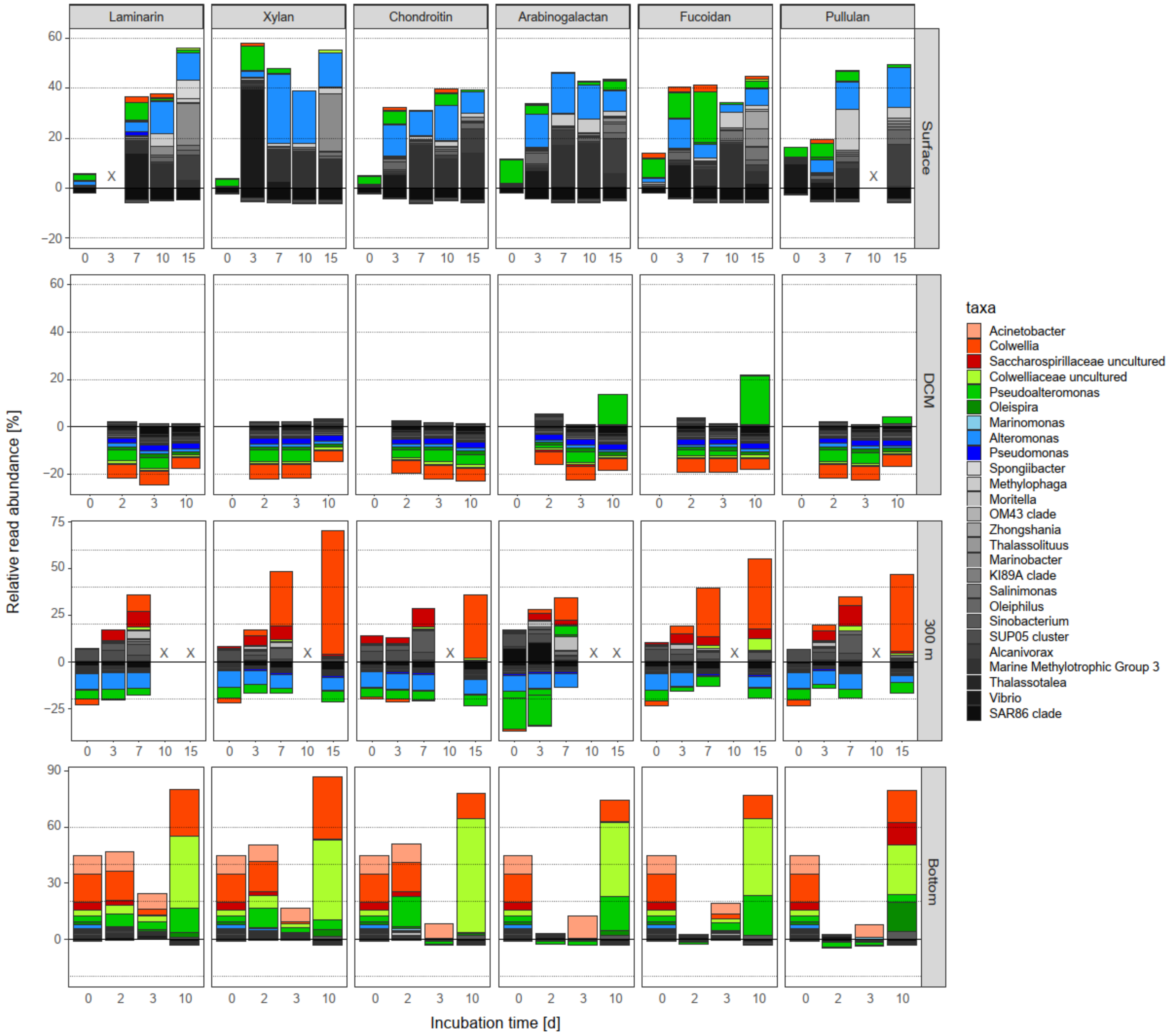


**Figure S9** Development of individual gammaproteobacterial taxa in FLA-PS amended incubations throughout the water column at Station 19. Bars represent the shift in 16S rRNA read abundance of individual Gammaproteobacteria relative to the initial bacterial community.

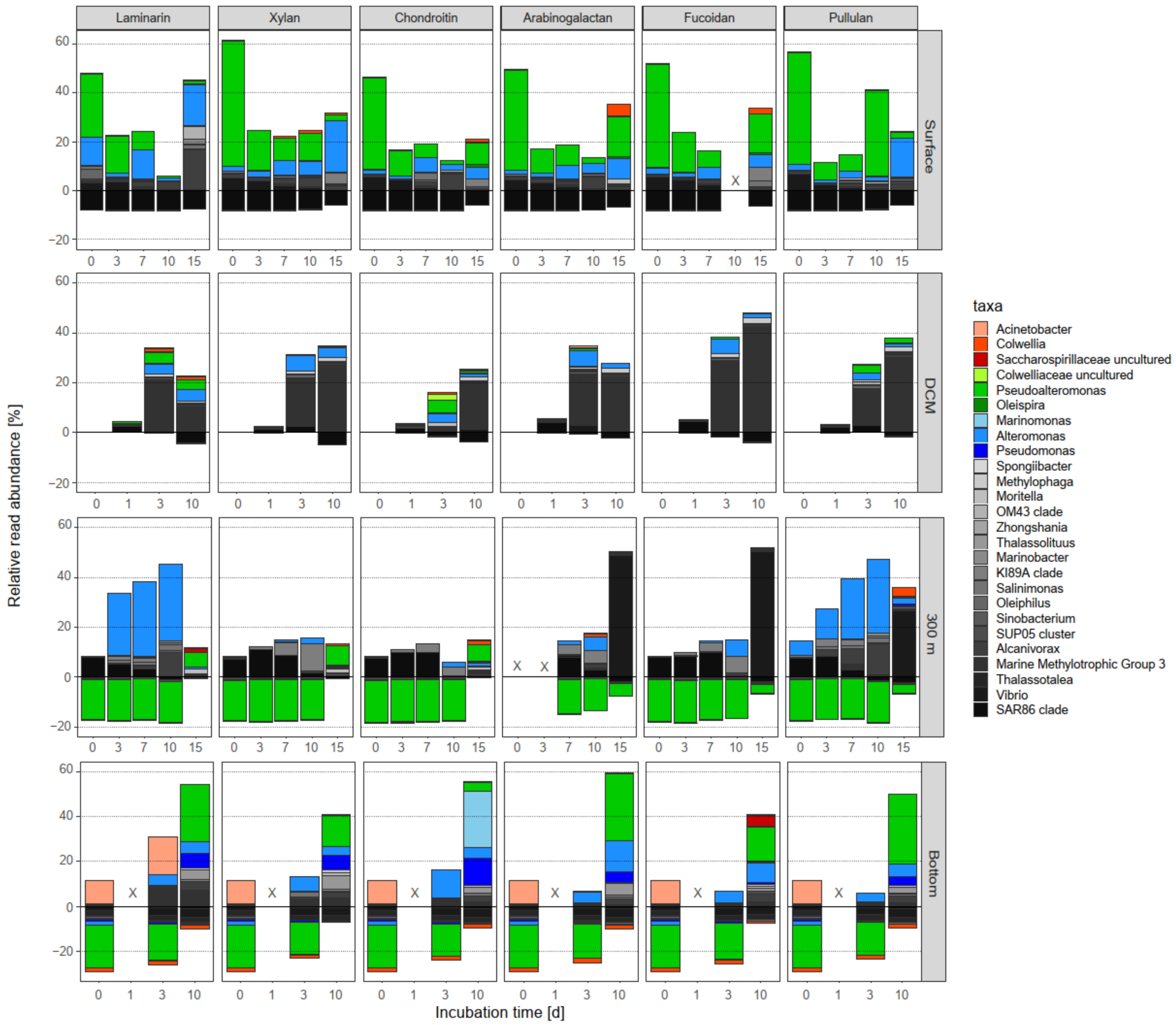


**Figure S10** Development of individual gammaproteobacterial taxa in FLA-PS amended incubations throughout the water column at Station 20. Bars represent the shift in 16S rRNA read abundance of individual Gammaproteobacteria relative to the initial bacterial community.

**Table S1** Bacterial protein production, initial bacterial abundance, and cell-specific protein production at Stations 18, 19, and 20.

| Station | **Station 18** | | | | **Station 19** | | | | **Station 20** | | | |
|---|---|---|---|---|---|---|---|---|---|---|---|---|
| Depth (m) | 2.5 | 104 | 300 | 3,190 | 6 | 33 | 318 | 4,325 | 5 | 64 | 300 | 5,580 |
| Bacterial Productivity (pmol $L^{-1}$ $h^{-1}$) | 6.04 ± 0.83 | 5.03 ± 1.20 | 0.46 ± 0.29 | 0.09 ± 0.08 | 88.1 ± 4.0 | 60.2 ± 7.9 | 1.2 ± 0.3 | 0.2 ± 0.1 | 191 ± 4.6 | 99.7 ± 7.1 | 23.5 ± 18.4 | 1.2 ± 0.1 |
| Initial bact. Abundance (cells $mL^{-1}$) | 4.00E+05 | 4.95E+05 | 1.43E+05 | 3.94E+04 | 9.71E+05 | 1.92E+06 | 1.44E+05 | 5.93E+04 | 3.66E+05 | 5.19E+05 | 2.45E+05 | 5.72E+03 |
| Cell-specific production (pmol $cell^{-1}$ $h^{-1}$) | 1.51E-08 | 1.02E-08 | 3.22E-09 | 2.28E-09 | 9.07E-08 | 3.14E-08 | 8.33E-09 | 3.37E-09 | 5.22E-07 | 1.92E-07 | 9.59E-08 | 2.10E-07 |
